# Stimulation of NCAM1-14.3.3.ζδ-derived Peptide Interaction Fuels Angiogenesis and Osteogenesis in Ageing

**DOI:** 10.1101/2024.01.16.575939

**Authors:** Taha Kadir Yesin, Hanyu Liu, Zhangfan Ding, Amit Singh, Qi Tian, Yuheng Zhang, Biswajyoti Borah, Junyu Chen, Anjali P. Kusumbe

**Author notes:** Author for correspondence: Anjali P. Kusumbe, MRC Weatherall Institute of Molecular Medicine, University of Oxford, John Radcliffe Hospital Oxford OX3 9DS, UK. These authors contributed equally (Taha Kadir Yesin, Hanyu Liu, Zhangfan Ding, Amit Singh).

## Abstract

The skeletal structure and bone marrow endothelium collectively form a critical functional unit essential for bone development, health, and aging. At the core of osteogenesis and bone formation lies the dynamic process of angiogenesis. In this study, we reveal a potent new endogenous anabolic NCAM1-14.3.3.ζδ-derived-Peptide interaction, which stimulates bone angiogenesis and osteogenesis during homeostasis, aging, and age-related bone diseases. Employing high-resolution imaging and inducible cell-specific mouse genetics, our results elucidate the pivotal role of the NCAM1-14.3.3.ζδ-derived-Peptide interaction in driving the expansion of Clec14a+ angiogenic endothelial cells. Notably, Clec14a+ endothelial cells express key osteogenic factors. The NCAM1-14.3.3.ζδ-derived-Peptide interaction in osteoblasts drives osteoblast differentiation, ultimately contributing to the genesis of new bone. Moreover, the NCAM1-14.3.3.ζδ-derived-Peptide interaction leads to a reduction in bone resorption. In age-associated vascular and bone loss diseases, stimulating the NCAM1-14.3.3.ζδ-derived-Peptide interaction not only promotes angiogenesis but also reverses bone loss. Consequently, harnessing the endogenous anabolic potential of the NCAM1-14.3.3.ζδ-derived-Peptide interaction emerges as a promising therapeutic modality for managing age-related bone diseases.

## Introduction

The skeleton and the bone marrow endothelium form a functional unit with great relevance in development, health, and ageing (Kusumbe et al., 2014; Salhotra & Shah, 2020). The bone marrow endothelium plays a central role in the maintenance of microenvironments required for regulating osteogenesis and haematopoiesis (Johnson et al., 2020; Owen-Woods & Kusumbe, 2022). Apart from supplying nutrients, the vasculature provides several inductive signals and secretory factors, so-called angiocrine signals/factors to regulate the tissue-specific microenvironments (Singh et al., 2019; Sivan et al., 2019).

Vasculature plays a key role in maintaining skeletal health and a dysregulation of the vasculature is speculated for several bone diseases like osteoporosis and osteoarthritis (Chen et al., 2021; Kumar et al., 2021; Stucker et al., 2020). Musculoskeletal disorders affect almost one in every two individuals posing a major health, psychological and economic burden on people, health systems and the Government (Lewis et al., 2019). With increasing life expectancy, ageing populations are more severely affected with osteoporosis and fracture (Brown et al., 2021; Karademir et al., 2015; Salari, Darvishi, et al., 2021; Salari, Ghasemi, et al., 2021; Xiao et al., 2022). Globally, osteoporosis is the most prevalent bone disease with a significant impact on health systems (Sözen et al., 2017; Sarafrazi et al., 2021). Existing therapies have mainly focused on slowing the rate of bone damage, with only a few drugs capable of promoting bone repair approved (Chindamo & Sapino, 2020; Liang & Burley, 2022). Due to issues like poor patient response, drug compliance, and drug-induced microfractures leading to atypical femur fractures (Tile & Cheung, 2020; Xiao et al., 2023), there is a pressing need for new therapies that effectively promote bone repair and regeneration in patients with musculoskeletal diseases, restoring tissue homeostasis and functional integrity.

Clinical management of non-healing fractures, which are characterized by delayed angiogenesis remains challenging (Schlundt et al., 2018; Stewart, 2019). In addition to the bone loss and fracture risk associated with cancer therapy; in bone marrow transplantations, metastatic bone disease, and cancer settings, the re-establishment of bone marrow microenvironments after chemotherapy and radiation is crucial for the effective treatment outcome and clinical management of these conditions (Chen et al., 2012; D’Oronzo et al., 2019; Singh et al., 2019; Tsukamoto et al., 2021; Vi et al., 2018; Wu et al., 2018). The bone vasculature and perivascular mesenchymal stem cells, which can differentiate to generate bone cells, adipocytes and chondrocytes, are essential for bone growth, and hematopoietic stem cell (HSC) maintenance (Hsu et al., 2021; James & Péault, 2019; Kusumbe et al., 2016). The manipulation of the vasculature is sufficient for the improvement of vascular HSC niche function, which suggests the existence of molecular pathways coupling the behaviour of endothelial cells and perivascular mesenchymal stem progenitor cells (MSPCs) in bone (Hanoun et al., 2014; Ramasamy, 2017). Thus, the prospect manipulating vasculature has tremendous potential to advance therapeutic interventions for the management of bone diseases and promote bone formation.

Notably, two members of the 14-3-3 family of adaptor proteins, 14-3-3β and 14-3-3ξ, differentially regulate osteoblasts (Pennington et al., 2018; Yasuda & Fukusumi, 2023). A bioactive endogenous 14.3.3.ζδ-derived-Peptide cleaved from 14-3-3ξ protein has been identified, influencing endothelium dependent monocyte migration during inflammation (Alassiri & Al Sufiani, 2023; Chimen et al., 2015). However, the role of this 14.3.3.ζδ-derived-Peptide remains unknown in bone homeostasis and ageing. Here, in this study using mouse genetics and high-resolution imaging, we unveil a new potent endogenous anabolic NCAM1-14.3.3.ζδ-derived-Peptide interaction in fuelling expansion of highly angiogenic Clec14a+ endothelial cells and promoting generation of new bone during homeostasis, aging, and age-related bone diseases.

## RESULTS

### Endogenous 14.3.3.ζδ-derived-Peptide Promoted Angiogenesis and Osteogenesis in Homeostasis and Ageing

To investigate the intrinsic regulatory role of 14.3.3.ζδ-derived-Peptide in bone remodelling under homeostatic conditions, we injected 14.3.3.ζδ-derived-Peptide in the mice (Figure 1A). 14.3.3.ζδ-derived-Peptide, when administered to adult mice, demonstrated a compelling role in regulating bone remodelling under normal physiological conditions. Our investigation involved a 10-day treatment period, during which 14.3.3.ζδ-derived-Peptide exhibited a remarkable ability to enhance both bone formation and bone mass (Figure 1B). Notably, 14.3.3.ζδ-derived-Peptide administration led to a significant increase in bone volume (BV/TV) and cortical thickness (Ct. Th) (Figure 1C). Importantly, a control peptide with an alternative 14-amino acid sequence derived from 14-3-3ξ, the parent protein of 14.3.3.ζδ-derived-Peptide, failed to elicit any discernible effects (Figure 1—figure supplement 1A).

**Figure 1.**
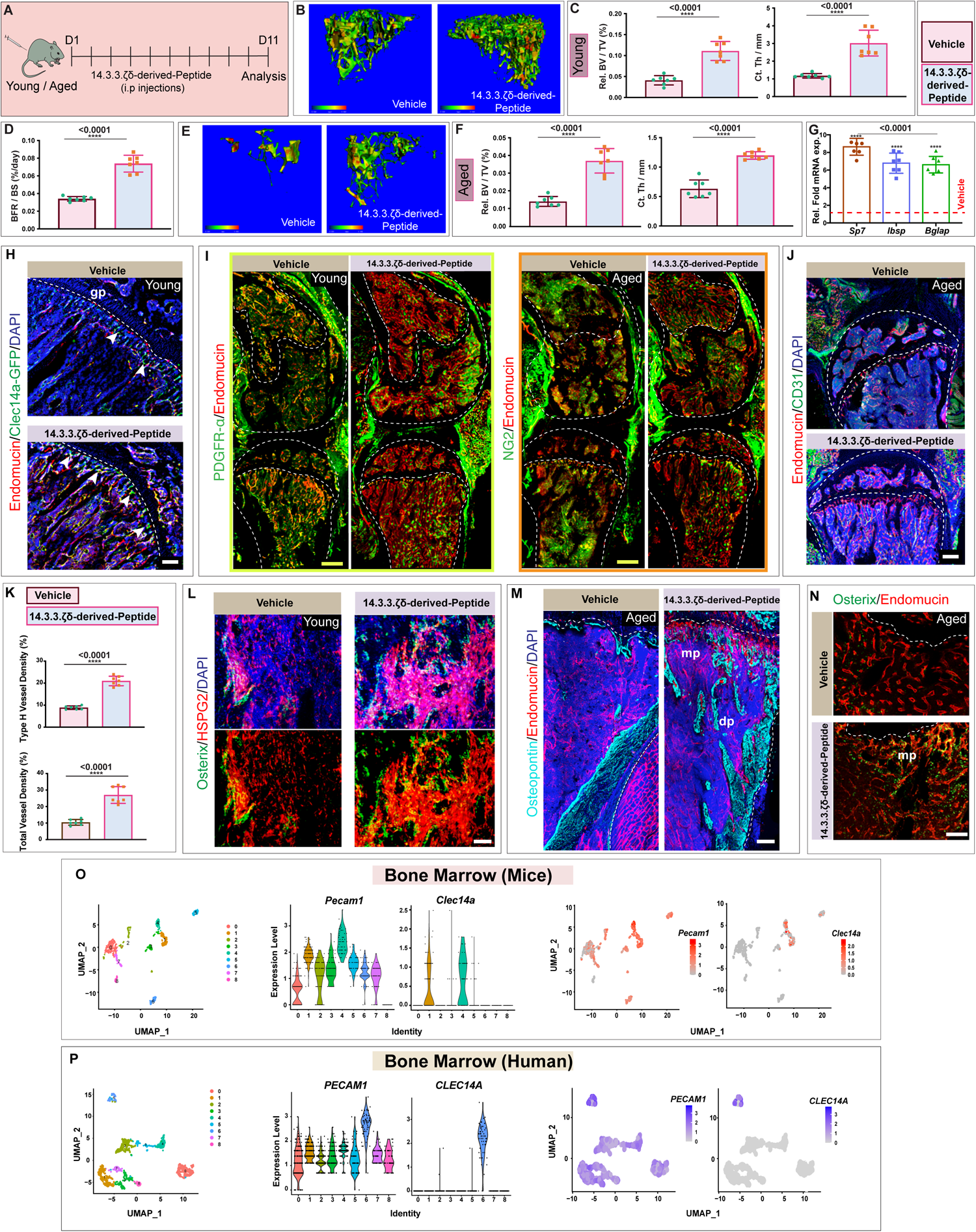
14.3.3.ζδ-derived-Peptide induces angiogenesis and osteogenesis in homeostasis and ageing. **(A)** Schematic depicting the experimental design, whereby young and aged mice were subjected to 10 day vehicle control (PBS) or 14.3.3.ζδ-derived-Peptide treatment prior to bone collection. **(B)** Representative (μ)-CT images of tibial bones from young mice treated with 14.3.3.ζδ-derived-Peptide compared to Vehicle mice **(C)** Quantitative analysis of relative bone volume (Rel. BV/TV, left panel) and cortical thickness (Ct. Th, right panel) of tibial bones from young mice treated with 14.3.3.ζδ-derived-Peptide compared to Vehicle (*n* =7). **(D)** Bone Formation Rate per Bone Surface (BFR/BS) in 14.3.3.ζδ-derived-Peptide treated aged mice compared to Vehicle (*n* =7). **(E)** Representative (μ)-CT images of tibial bones from aged mice treated with 14.3.3.ζδ-derived-Peptide compared to Vehicle. **(F)** Quantitative analysis of relative bone volume (Rel. BV/TV, left panel) and cortical thickness (Ct. Th, right panel) of tibial bones from aged mice treated with 14.3.3.ζδ-derived-Peptide compared to Vehicle. (*n* =7). **(G)** Relative fold *mRNA* expression of *Sp7, Ibsp, and Bglap* in aged mice treated with 14.3.3.ζδ-derived-Peptide compared to vehicle (*n* =7). **(H)** 3D images of tibial bone from young mice treated with 14.3.3.ζδ-derived-Peptide compared to Vehicle; showing Endomucin (red), Clec14a-GFP (green), and DAPI (blue). **(I)** 3D images of bones from young mice labeled with PDGFR-α (green) and Endomucin (red) (left panels); aged mice labeled with NG2 (green) and Endomucin (red) (right panels). Both sets of images compare mice treated with 14.3.3.ζδ-derived-Peptide to Vehicle. **(J)** 3D images of tibial bone from aged mice treated with 14.3.3.ζδ-derived-Peptide compared to Vehicle; showing Endomucin (red), CD31 (green), and DAPI (blue). **(K)** Quantification of type H vessel and total vessel density in tibial bones from Vehicle and 14.3.3.ζδ-derived-Peptide treated mice (*n* =7). **(L)** 3D images of tibial bone from young mice treated with 14.3.3.ζδ-derived-Peptide compared to Vehicle; showing immunolabelling for Osterix (green), Fibronectin (red), and DAPI (Blue). **(M)** 3D images of tibial bone from aged mice treated with 14.3.3.ζδ-derived-Peptide compared to Vehicle; showing Osteopontin (Cyan), Endomucin (red), and DAPI (blue). **(N)** 3D images of tibial bone from aged mice treated with 14.3.3.ζδ-derived-Peptide compared to Vehicle; showing Osterix (green) and Endomucin (red). **(O)** UMAP projections and Violin Plots of single cells from mice bone marrow showing *Pecam1* and *Clec14a* expressions in different endothelial cell subsets. **(P)** UMAP projections and Violin Plots of single cells from human bone marrow showing *PECAM1* and *CLEC14A* expressions in different endothelial cell subsets. **Data Information:** Two-tailed unpaired t tests (C,D,F,G and K). nsp > 0.05, **p < 0.01, ***p < 0.001, and ****p < 0.0001. Scale bars: white, 50 μm; blue 200 μm; yellow, 300 μm. n represents biological replicates. Growth plate: gp; Diaphysis: dp; Metaphysis: mp.

Recognizing that aging is associated with bone loss and bone-related disorders, we extended our investigation to aged mice. Remarkably, a 10-day treatment with 14.3.3.ζδ-derived-Peptide proved sufficient to induce a substantial increase in bone mass, accompanied by elevated expression of *Sp7, Ibsp and Bglap* genes associated with osteogenesis in the aged mice (Figures 1D-G).

Moreover, administration of 14.3.3.ζδ-derived-Peptide; both in young and aged mice, led to increased significant augmentation of angiogenesis, a process crucial for vascular growth and expansion. Both young and aged mice exhibited increased expression of vascular markers and increased vascular density, validated through imaging thick bone slices and quantitative analysis (Figures 1H-K). This augmented angiogenic response correlated closely with an increased accumulation of Clecl14a^+^ cells near the growth plate (Figure 1H) - a notable marker expressed by highly angiogenic endothelial cells with a pivotal role in embryonic and pathological angiogenesis, as well as adhesion and migration (Khan et al., 2017; Kim et al., 2018; Noy et al., 2015). Recent studies have also emphasized the function of Group 14 C-type lectins, of which Clec14a is a member (Khan et al., 2019).

The observed structural changes and increased bone mass in 14.3.3.ζδ-derived-Peptide-treated mice were characterized by an increased abundance of Osterix-expressing osteoblasts and Osteopontin expression (Figures 1L-N). However, no changes were observed in number of adipocytes as analysed by Perilipin immunostaining (Figure 1—figure supplement 1B). Taken together, various vascular and osteoblast cell markers demonstrated a remarkable increase following 14.3.3.ζδ-derived-Peptide treatment in both young and aged animals (Figures 1L-N).

In summary, a 10-day 14.3.3.ζδ-derived-Peptide treatment unequivocally induced bone formation and boosted bone mass in both young and aged animals. Notably, the effectiveness of 14.3.3.ζδ-derived-Peptide in promoting bone formation, as indicated by its impact on BV/TV after 10 days, rivals that achieved by established bone-targeting drugs such as zoledronic acid (over 3 weeks) or parathyroid hormone (over 4 weeks) (Zhu et al., 2014; Zweifler & Koh, 2021). This suggests that 14.3.3.ζδ-derived-Peptide exhibits comparable efficacy to current standard-of-care treatments in inducing bone formation. These comprehensive findings, presented for the first time, unveil the pro-osteogenic actions of 14.3.3.ζδ-derived-Peptide under homeostatic conditions without inflammatory stimuli.

### 14.3.3.ζδ-derived-Peptide Treatment induced Expansion of Clec14a+ Angiogenic Endothelial Cells

After 14.3.3.ζδ-derived-Peptide treatment, our results clearly demonstrated the expansion of Clec14a+ endothelial cells located near the growth plate. Moreover, these Clec14a+ endothelial cells were detected both in mouse and human bone marrow as revealed by the single-cell sequencing analysis (Figures 1O and P). Clec14a, or C-type lectin domain family 14 member A, is a gene that codes for a protein with a C-type lectin domain. C-type lectins are a family of proteins with carbohydrate-binding domains, and they often play roles in immune response and cell adhesion. As stated above Clec14a expressing cells have been found in tumour endothelium and expression of Clec14a is associated with high angiogenic activity (Mancuso et al., 2014). Therefore, to gain a deeper understanding of the role of these cells in bone development, especially during the 14.3.3.ζδ-derived-Peptide-induced increase in bone mass, we conducted genetic lineage tracing of these cells. To achieve this, we generated a tamoxifen-inducible *Clec14a Cre ER^T2^*line in *Clec14a PiggyBac-on-BAC* Mouse Transgenics (Figure 2—figure supplements 1A-D). The utilization of genetic lineage tracing, a powerful investigative tool, enabled us to explore processes such as cell amplification, differentiation, and migration. Employing *Clec14a-CreER^T2^ x ROSA26-mT/mG* double transgenics and administering tamoxifen, we identified Clec14a+ cells strategically positioned near the growth plate, at the angiogenic forefront of the bones in both juvenile and adult mice. In the juvenile mice, these cells were abundant in the metaphysis region of the bones. Notably, these cells exhibited high expression of type H endothelium markers such as Endomucin (Figure 2A).

**Figure 2.**
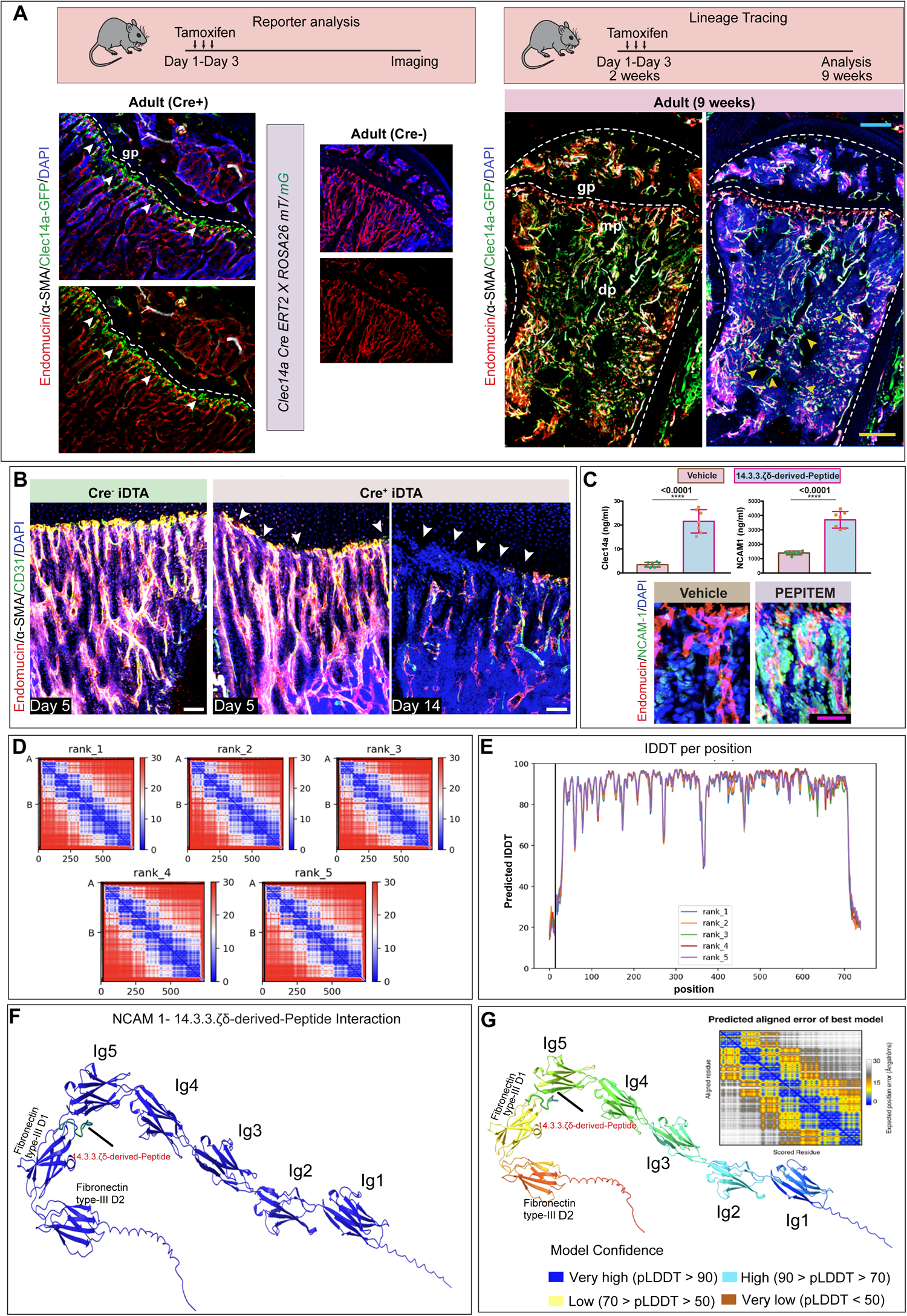
Lineage tracing of Celec14a+ endothelial cells and 3D modelling of NCAM1-14.3.3.ζδ-derived-Peptide interactions. **(A)** Schematic depicting the experimental design, whereby *Clec14a Cre ER^T2^ x Rosa26 - mT/mG* mice were generated. Tibial bones were collected for analysis. 3D images showing Endomucin (red), α-SMA (White), Clec14a-GFP (Green) and DAPI (blue); CD31 (red), and DAPI (blue). **(B)** 3D images from Cre^-^ iDTA or Cre+ iDTA, *Clec14a Cre ER^T2^ X iDTA* mice were analyzed at 5- and 14-days after the last dose of tamoxifen; labeled with CD31 (green), Endomucin (red), α-SMA (white) and DAPI (blue). Arrowheads indicate loss of endothelial cells near the growth plate region. **(C)** ELISA quantification of NCAM1 and Clec14a concentration from 14.3.3.ζδ-derived-Peptide treated mice compared to Vehicle. (n = 7). 3D images of tibial bone from mice treated with 14.3.3.ζδ-derived-Peptide compared to Vehicle; showing Endomucin (red), NCAM-1 (green), and DAPI (blue). **(D)** Predicted aligned error (PAE). Five different models predicted aligned error (PAE) obtained from AlphaFold. The colour code denotes the anticipated positional discrepancy at residue x under the condition that the predicted and actual structures align with y and are measured in Ångströms. **(E)** The local distance difference test (IDDT) of the best-predicted model **(F)** NCAM 1-14.3.3.ζδ-derived-Peptide Interaction complex prediction. The blue colour indicates NCAM 1 protein, and the green colour indicates 14.3.3.ζδ-derived-Peptide. IG1 domain contain (20-111), IG2 domain (116-189), IG3 domain (212-30), IG4 domain (308-403), IG5 domain (406-191), Fibronectin type -III domain 1(499-598), Fibronectin type -III domain 2(600-695) amino acid. **(G)** Best predicted model of NCAM 1-14.3.3.ζδ-derived-Peptide Interaction. The colour indicates model confidence. **Data Information:** Two-tailed unpaired t tests (C). nsp > 0.05, **p < 0.01, ***p < 0.001, and ****p < 0.0001. Scale bars: magenda 25 μm; white, 50 μm; yellow, 300 μm. n represents biological replicates. Growth plate: gp; Diaphysis: dp; Metaphysis: mp.

Further genetic lineage tracing of these cells using *Clec14a-CreER^T2^ x ROSA26-mT/mG* double transgenics towards which tamoxifen administration was performed in 3-weeks-old mice, followed by analysis in adult 9-weeks-old mice, revealed that these Clec14a+ cells hierarchically positioned upstream of other endothelial cells (Figure 2A). Clec14a+ cells which expressed the markers for type H endothelium gave rise to and differentiated into various endothelial cell types in adult bones.

To further investigate the impact of Clec14a angiogenic cells on bone formation during 14.3.3.ζδ-derived-Peptide treatment, we implemented a targeted strategy aimed at directly affecting and depleting these cells. To selectively deplete the Clec14a+ cell population and assess their influence on bone formation during 14.3.3.ζδ-derived-Peptide treatment, we utilized the *Clec14a-Cre ER^T2^* mouse line crossed with mice expressing a tamoxifen-inducible diphtheria toxin antigen (iDTA) (Figure 2—figure supplement 1F). Tamoxifen injections were administered to both Cre- and Cre+ mice. As expected, Cre+ iDTA mice exhibited suppressed expansion of Clec14a cells and led to reduced expression of Clec14a expressing cells. Moreover, analysis of the Cre+ iDTA mice demonstrated loss of endothelial cells near the growth plate region (Figure 2B). Moreover, analysis of Clec14a+ endothelial cells had high expression of several osteogenic factors compared to Clec14a-endothelial cell in bones (Figure 2—figure supplement 1E).

Microcomputed tomography (m-CT) analyses of Cre+ iDTA mice, 14.3.3.ζδ-derived-Peptide treatment demonstrated that the removal of Clec14a+ cells did not lead to an increase in bone mass following 14.3.3.ζδ-derived-Peptide treatment (Figure 2—figure supplement 1F). These collective findings strongly support the crucial role of Clec14a+ cells in facilitating bone formation during 14.3.3.ζδ-derived-Peptide treatment. The targeted depletion of these cells during 14.3.3.ζδ-derived-Peptide treatment not only influenced the expression of key osteogenic markers but also resulted in a noticeable decrease in overall bone density, as confirmed by micro-CT assessments. These results highlight the interplay between Clec14a angiogenic cells and the regulatory mechanisms governing bone homeostasis in the context of 14.3.3.ζδ-derived-Peptide treatment.

### Anabolic NCAM-1-14.3.3.ζδ-derived-Peptide Interaction drives Angiogenesis and Bone Formation

In our investigation into the osteogenic actions of 14.3.3.ζδ-derived-Peptide, we sought to identify the receptor responsible for its effects on endothelial cells. Despite the known presence of CDH15 in primary endothelial cells (Matsubara et al., 2020), treatment with a CDH15 agonistic antibody failed to impact bone formation and bone mass during 14.3.3.ζδ-derived-Peptide treatment, decisively ruling out CDH15 as the mediator of 14.3.3.ζδ-derived-Peptide’s (Figure 3—figure supplement 1A). These compelling findings strongly suggest that 14.3.3.ζδ-derived-Peptide’s osteogenic mechanism is novel and separate from its well-established immunopeptide actions in inflammatory challenges.

Drawing parallels with angiogenesis marker Clec14a which was increased after 14.3.3.ζδ-derived-Peptide treatment, we identified NCAM-1 which is a known marker for highly angiogenic cells and associated with tumour endothelium (Zhang et al., 2019), and immunostaining and ELISA analysis confirmed its expression in endothelial cells after 14.3.3.ζδ-derived-Peptide treatment (Figure 2C). To investigate whether NCAM-1 serves as a receptor for 14.3.3.ζδ-derived-Peptide, we utilized the AlphaFold2_Multimer neural network in conjunction with ChimeraX to generate five 3D models predicting the binding and interaction of 14.3.3.ζδ-derived-Peptide with NCAM-1 (Figures 2D-G). NCAM-1 structurally comprises 5 IgG-like domains and 2 fibronectin-III-like domains. The protein sequences of Neural Cell Adhesion Molecule 1 (NCAM-1, obtained from Uniprot with accession code H7BYX6) and 14.3.3.ζδ-derived-Peptide (SVTEQGAELSNEER) were inputted into AlphaFold2_multimer, a tool run on Google Colab, resulting in the production of five distinct models (Figure 2D). The selection of the most reliable models was based on the predicted local distance difference test (pLDDT), prioritizing those with the highest predicted confidence (Figure 2E). Subsequently, ChimeraX was employed for constructing a 3D model that illustrates the interaction between NCAM-1 and 14.3.3.ζδ-derived-Peptide (Figures 2F and G). These predictive 3-D models suggest that 14.3.3.ζδ-derived-Peptide mediates its effects via the NCAM-1. Notably, our analysis unveiled a substantial increase in *Ncam1* expression on both osteoblasts and Clec14a+ endothelial cells after 14.3.3.ζδ-derived-Peptide treatment (Figure 3— figure supplement 1B), aligning with previous evidence implicating NCAM-1 in the regulation of osteoblast differentiation through Wnt/β-catenin and PI3K-AKT signalling pathways (Agas & Hanna, 2021; Huang et al., 2021; Lara-Castillo et al., 2023). Therefore, next we interrogated the role of NCAM1 *in vivo* in mouse bones. Function-blocking experiments with an NCAM-1 antibody demonstrated that inhibiting NCAM-1 abolished the osteogenic effect of 14.3.3.ζδ-derived-Peptide treatment (Figure 3— figure supplement 1C).

**Figure 3.**
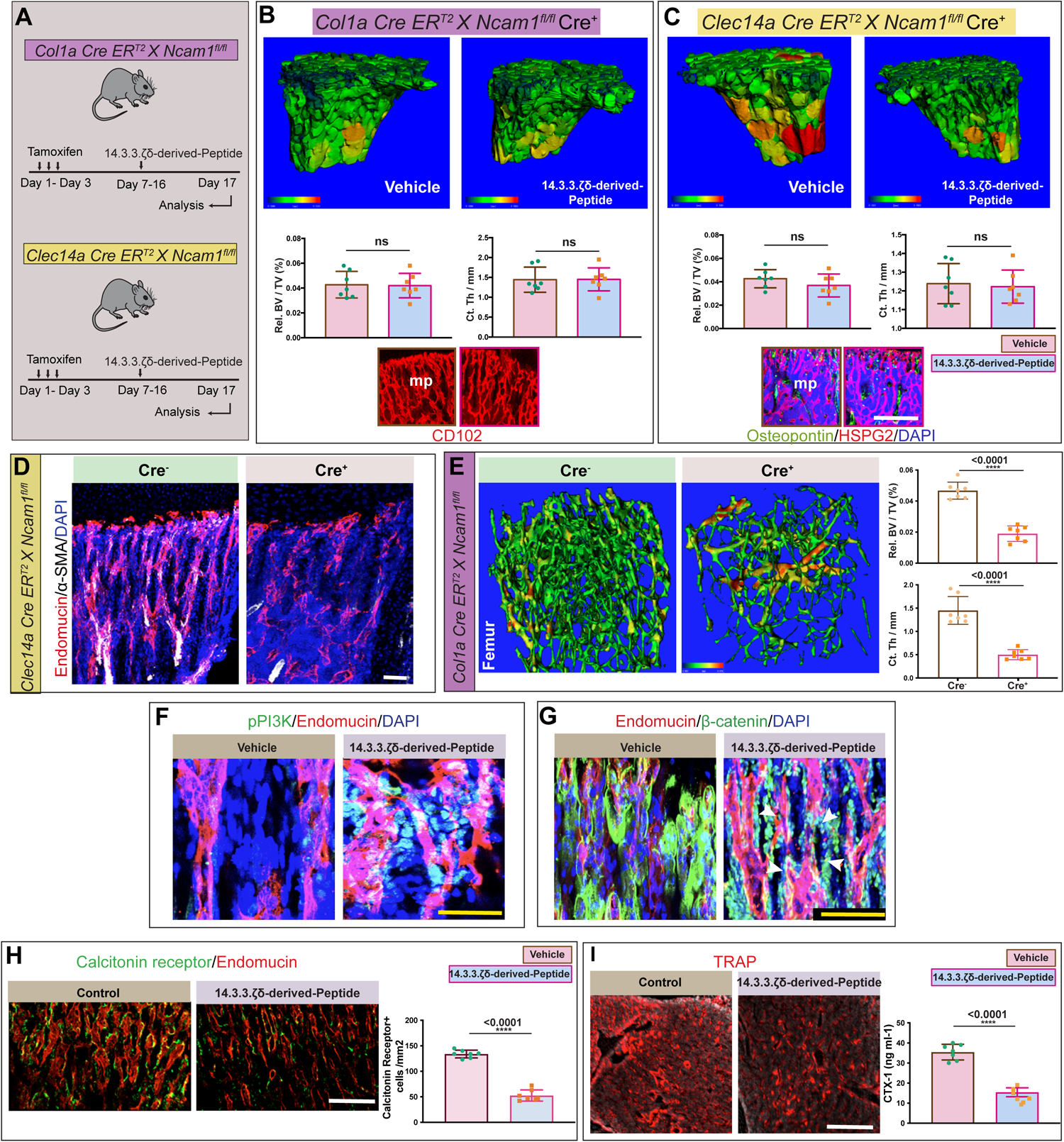
NCAM1 regulates 14.3.3.ζδ-derived-Peptide induced angiogenesis and osteogenesis. **(A)** Schematic depicting the experimental design, whereby tamoxifen-inducible *Col1a Cre ER^T2^ X Ncam1^fl/fl^* and *Clec14a Cre ER^T2^ X Ncam1^fl/fl^* mice untreated or subjected to 14.3.3.ζδ-derived-Peptide post-tamoxifen administration, and bones were subsequently collected and stained for imaging. **(B)** Representative (μ)-CT images of tibiae from *Col1a Cre ER^T2^ X Ncam1^fl/fl^*, Cre+ mice treated with 14.3.3.ζδ-derived-Peptide compared to Vehicle. Quantitative analysis of relative bone volume (Rel. BV/TV, left panel) and cortical thickness (Ct. Th, right panel) of the bones (n = 7). 3D images showing CD102 (red). **(C)** Representative (μ)-CT images of tibiae from *Clec14a Cre ER^T2^ X Ncam1^fl/fl^*, Cre+ mice treated with 14.3.3.ζδ-derived-Peptide compared to Vehicle. Quantitative analysis of relative bone volume (Rel. BV/TV, left panel) and cortical thickness (Ct. Th, right panel) of the bones (n = 7). 3D images showing Endomucin (red), HSPG2 (green) and DAPI (blue). **(D)** 3D images of tibial bone from Cre^-^ and Cre^+^, *Clec14a Cre ER^T2^ X Ncam1^fl/fl^* mice; showing Endomucin (red), α-SMA (White) and DAPI (blue). **(E)** Representative (μ)-CT images of femur from Cre^-^ and Cre^+^, *Col1a Cre ER^T2^ X Ncam1^fl/fl^* mice. Quantitative analysis of relative bone volume (Rel. BV/TV) and cortical thickness (Ct. Th) of the bones (n = 7). **(F)** 3D images of tibial bone from mice treated with 14.3.3.ζδ-derived-Peptide compared to Vehicle; showing pPI3K (green), Endomucin (red), and DAPI (blue). **(G)** 3D images of tibial bone from mice treated with 14.3.3.ζδ-derived-Peptide compared to Vehicle; showing Endomucin (red), β-catenin (green), and DAPI (blue). **(H)** 3D images of tibial bone from mice treated with 14.3.3.ζδ-derived-Peptide compared to Control mice; showing osteoclasts labeled with Calcitonin receptor (green) and Endomucin (red). Quantification of Calcitonin Receptor+ cells from 14.3.3.ζδ-derived-Peptide treated mice compared to Vehicle. (n = 7). **(I)** 3D images of tibial bone from mice treated with 14.3.3.ζδ-derived-Peptide compared to Control mice; showing osteoclasts labeled with TRAP (red). Quantification of CTX-1 levels from 14.3.3.ζδ-derived-Peptide treated mice compared to Vehicle. (n = 7). **Data Information:** Two-tailed unpaired t tests (B,C,E,H and I); nsp > 0.05, **p < 0.01, ***p < 0.001, and ****p < 0.0001. Scale bars: white, 50 μm; yellow, 300 μm. n represents biological replicates. Metaphysis: mp

Next, we interrogated the *in vivo* functional role of NCAM-1 during 14.3.3.ζδ-derived-Peptide treatment in endothelial cells and osteoblasts, we employed osteoblast-specific *Col1a Cre ER^T2^* and endothelial *Clec14a Cre ER^T2^* mouse models. Tamoxifen injections were administered to both Cre+ and Cre-adult mice utilizing osteoblast-specific *Col1a Cre ER^T2^ X Ncam1^fl/fl^* and endothelial-specific *Clec14a Cre ER^T2^ X Ncam1^fl/fl^* mice (Figure 3A).

Analysis of endothelial-specific and osteoblast-specific Ncam1-loss-of-function Cre+ mouse bones revealed that this specific loss of *Ncam1* in endothelial or osteoblast cells resulted in no increase in bone mass and bone density following 14.3.3.ζδ-derived-Peptide treatment (Figures 3B and C). Subsequently, we examined osteoblast-specific *Col1a Cre ERT2 X Ncam1^fl/fl^* and endothelial-specific *Clec14a Cre ERT2 X Ncam1^fl/fl^* mice during postnatal bone development after tamoxifen administration in 1-week-old mice without 14.3.3.ζδ-derived-Peptide treatment. Tamoxifen was administered to both Cre+ and Cre-mice (Figures 3D and E). *Col1a Cre ER^T2^ X Ncam1^fl/fl^* mice exhibited a notable reduction in bone mass and bone density (Figure 3E), aligning with prior studies highlighting the regulatory role of NCAM-1 in osteoblast differentiation (Cheng et al., 2021).

Analysis of *Clec14a Cre ER^T2^ X Ncam1^fl/fl^* mice demonstrated reduced vessel density in Cre+ mouse bones and diminished endothelial cell proliferation (Figure 3D), consistent with earlier studies implicating NCAM-1 in the regulation of endothelial cell proliferation through the PI3K-AKT signalling pathway (Xiang et al., 2021). Immunostaining confirmed increased levels of phosphorylated PI3K (pPIK3) (Figure 3F).

Prior research has suggested the involvement of NCAM-1 in the regulation of osteoblast differentiation through the Wnt/β-catenin pathway (Cheng et al., 2021). To explore this further, we assessed beta-catenin expression and localization. Immunostaining of bones post-14.3.3.ζδ-derived-Peptide treatment revealed a remarkable increase in intracellular β-catenin (Figure 3G).

In summary, these findings indicate that NCAM-1 plays a pivotal role in modulating bone mass, bone density, vessel density, and endothelial cell proliferation during 14.3.3.ζδ-derived-Peptide treatment *in vivo*. These effects involve PI3K-AKT signalling and the Wnt/β-catenin pathway, shedding light on the intricate mechanisms underlying NCAM-1’s functional significance in the context of 14.3.3.ζδ-derived-Peptide treatment. These comprehensive findings collectively unveil a novel and impactful mechanism by which 14.3.3.ζδ-derived-Peptide orchestrates β-catenin translocation in osteoblasts through its interaction with NCAM-1, providing valuable insights for potential therapeutic applications in bone-related disorders.

### NCAM1-14.3.3.ζδ-derived-Peptide Interaction Leads Suppression of Bone Resorption

Recognizing the intricate regulation of bone homeostasis by osteoblasts and osteoclasts (Kim et al., 2020), our investigation aimed to elucidate whether the observed increase in bone mass was a result of heightened osteoblast activity or diminished osteoclast function. The finely tuned interplay between osteoblasts and osteoclasts involves intricate regulation, with heightened osteoblast activity often matched by increased osteoclastogenesis to maintain balance (Kim et al., 2020; Marahleh et al., 2023). To comprehensively understand NCAM1-14.3.3.ζδ-derived-Peptide pathway impact on bone dynamics, we evaluated its effects on osteoclast function. Examination of 14.3.3.ζδ-derived-Peptide-treated mice unveiled a substantial reduction in osteoclast numbers compared to controls, as evidenced by diminished tartrate-resistant acid phosphatase (TRAP)-positive cells, Calcitonin Receptor positive cells and CTX-1 levels (Figures 3H and I). Next, we analysed of *Clec14a Cre ER^T2^ X Ncam1^fl/fl^* mice *and Col1a Cre ER^T2^ X Ncam1^fl/fl^* mice treated with 14.3.3.ζδ-derived-Peptide for osteoclast number and function. Examination of these mice demonstrated no changes in osteoclast numbers and function, as analysed by assessment of Calcitonin Receptor positive cells and CTX-1 levels (Figure 3—figure supplement 1D). These results demonstrates that NCAM1-14.3.3.ζδ-derived-Peptide interaction is required for reduction in osteoclast numbers and bone resorption.

### NCAM1-14.3.3.ζδ-derived-Peptide drives Angiogenesis and Reverses Bone Loss in Age-related Bone Diseases

In light of the substantial pro-angiogenic effects demonstrated by NCAM1-14.3.3.ζδ-derived-Peptide interaction, our investigation sought to comprehensively assess its therapeutic efficacy in treating musculoskeletal diseases associated with vascular and bone loss. Osteoporosis, a prevalent musculoskeletal condition linked to cancellous bone loss and an increased risk of fractures (Sözen et al., 2017; Sarafrazi et al., 2021), often lacks interventions targeting vascular and osteoblast-driven repair processes. NACM1-14.3.3.ζδ-derived-Peptide, with its remarkable pro-angiogenic and osteogenic effects, emerged as a compelling candidate for such intervention to stimulate angiogenesis and osteogenesis.

In a murine model of osteoporosis induced by ovariectomy, 14.3.3.ζδ-derived-Peptide therapy effectively reduced bone loss compared to the vehicle control (Figure 4A). Mice treated with 14.3.3.ζδ-derived-Peptide demonstrated increased expression of *Clec14a* and *Ncam1* maintained bone volume density (BV/TV) and cortical thickness levels comparable to the control group, in stark contrast to the significant reduction observed in the vehicle control group. Imaging for osteogenic markers demonstrated increased expression (Figure 4A, Figure 4—figure supplement 1). Importantly, 14.3.3.ζδ-derived-Peptide treatment in the *Clec14a Cre ER^T2^ X Ncam1^fl/fl^* Cre+ ovariectomized mice did not reverse the bone mass confirming that NCAM1-14.3.3.ζδ-derived-Peptide interaction in Clec14a+ endothelium is required to induce bone mass in 14.3.3.ζδ-derived-Peptide treated mice. (Figure 4A).

**Figure 4.**
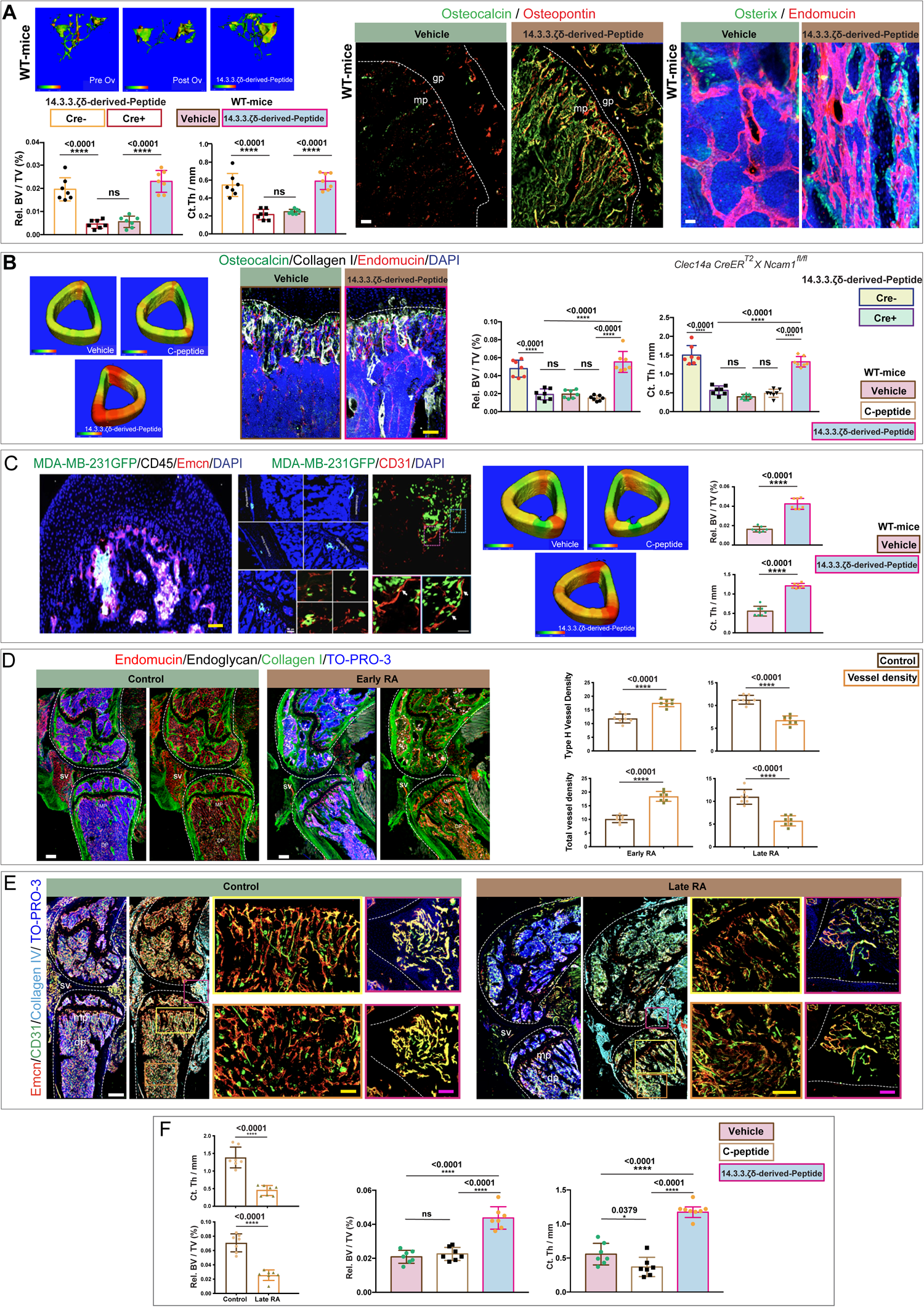
14.3.3.ζδ-derived-Peptide treatment inducedangiogenesis and bone formation in aged related musculoskeletal diseases. **(A)** Mouse model of osteoporosis. Representative (μ)-CT images of tibial bones from pre-ovariectomy, post ovariectomy and 14.3.3.ζδ-derived-Peptide treated mice. Quantitative analysis of relative bone volume (Rel. BV/TV, left panel) and cortical thickness (Ct. Th, right panel) of tibial bones from Cre- or Cre+, *Clec14a Cre ER^T2^ X Ncam1^fl/fl^* and Vehicle or 14.3.3.ζδ-derived-Peptide treated wild-type mice (*n* =7). 3D images of tibial bone from mice treated with 14.3.3.ζδ-derived-Peptide compared to Vehicle labeled with Ostecalcin (green) and Osteopontin (red) (left panel); Osterix (green) and Endomucin (red) (right panel). **(B)** Mouse model of osteoarthritis. Representative (μ)-CT images of tibial bones from Vehicle, C-peptide and 14.3.3.ζδ-derived-Peptide treated mice. 3D images of tibial bone from mice treated with 14.3.3.ζδ-derived-Peptide compared to Vehicle; showing Osteocalcin (green), Collagen I (White), Endomucin (red) and DAPI (blue). Quantitative analysis of relative bone volume (Rel. BV, left panel) and cortical thickness (Ct. Th, right panel) of tibial bones from Cre- or Cre+, *Clec14a Cre ER^T2^ X Ncam1^fl/fl^* and vehicle, C-peptide or 14.3.3.ζδ-derived-Peptide treated wild-type mice (n=7). **(C)** Mouse model of bone metastasis. 3D images from the tibial bone of mice that develop metastases, taken after intracardiac injection of tumor cells; showing localization of MDA-MB-231 tumor cells, labeled with GFP (green), CD45 (white), Endomucin (red) and DAPI (blue) (left panel); GFP (green), CD31 (red) and DAPI (blue) (right panel). Note the localization of these cells in the metaphysis region near the growth plate. Representative (μ)-CT images of tibial bones from Vehicle, C-peptide and 14.3.3.ζδ-derived-Peptide treated mice. Quantitative analysis of relative bone volume (Rel. BV/TV) and cortical thickness (Ct. Th) of tibial bones (n=7). **(D)** Mouse model of Rheumatoid Arthritis (RA). 3D images of bones from RA early compared to control mice; labeled with Endomucin (red), Endoglycan (white), Collagen I (green) and TO-PRO3 (blue). Quantification of type H and total vessel density in tibial bones from RA early and late mice models (n=7). **(E)** 3D images of bones from RA late mouse model compared to control; labeled with Endomucin (red), CD31 (green), Collagen IV (cyan) and TO-PRO3 (blue). **(F)** Quantitative analysis of relative bone volume (Rel. BV/TV) and cortical thickness (Ct. Th) – control and late RA (left panel) and vehicle, C-peptide and 14.3.3.ζδ-derived-Peptide treated (right panel) (n=7). **Data Information:** Two-tailed unpaired t tests (C, D and F); One-way ANOVA tests with Tukey’s multiple comparisons tests (A,B and F). nsp > 0.05, **p < 0.01, ***p < 0.001, and ****p < 0.0001. Scale bars: magenda 25 μm; white, 50 μm; yellow, 300 μm. n represents biological replicates. Synovial membrane Volume; sv; Growth plate: gp; Diaphysis: dp; Metaphysis: mp.

**Figure 5.**
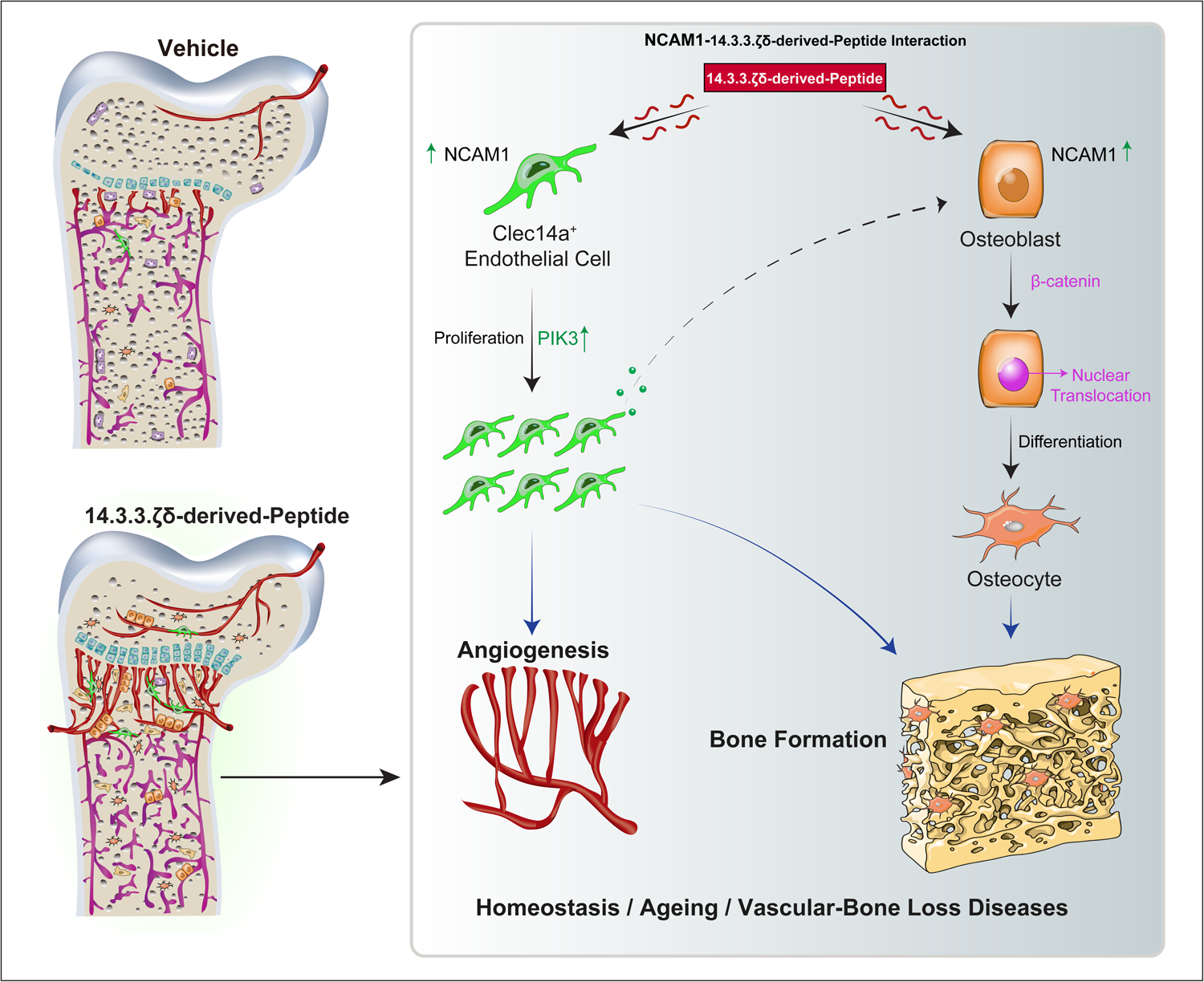
Schematic showing 14.3.3.ζδ-derived-Peptide induced angiogenesis and osteogenesis through expansion of CLEC14+ cells and NCAM1 interactions.

Subsequently, we investigated the potential of NACM1-14.3.3.ζδ-derived-Peptide interaction in a mouse model of osteoarthritis, a bone loss disease. 14.3.3.ζδ-derived-Peptide treatment reversed bone loss, induced angiogenesis as analysed by upregulation of the expression of *Clec14a* and *Ncam1* as evidenced by micro-CT analysis and immunostaining for osteogenic markers (Figures 4B, Figure 4—figure supplement 1). Once again, 14.3.3.ζδ-derived-Peptide treatment in the *Clec14a Cre ER^T2^ X Ncam1^fl/fl^* Cre+ mice did not reverse the bone mass confirming that NCAM1-14.3.3.ζδ-derived-Peptide interaction in Clec14a+ endothelium is required to induce bone mass in 14.3.3.ζδ-derived-Peptide treated mice with osteoarthritis. (Figure 4B). Further exploration of NACM1-14.3.3.ζδ-derived-Peptide interaction extended to bone metastasis, where treatment with 14.3.3.ζδ-derived-Peptide reversed bone loss in the bone metastasis mouse model (Figure 4C). Analysis of a rheumatoid arthritis model, characterized by bone loss and decreased vascular density, revealed a significant loss of vessel density in advanced disease stages (Figures 4D and E). This observation contrasted with the reported increase in angiogenesis and endothelial cells during inflammation (Hellenthal & Brabenec, 2022), suggesting a coupled relationship between angiogenesis and osteogenesis even in inflammatory settings. The loss of vasculature in advanced disease as compared to early stage was associated with a subsequent loss of bone mass (Figures 4D and E). Treatment with 14.3.3.ζδ-derived-Peptide led to an increase in angiogenesis as assessed by expression of *Clec14a* and *Ncam1* expressions in bone mass and cortical bone thickness (Figure 4F, Figure 4— figure supplement 1).

In summary, these findings position NCAM1-14.3.3.ζδ-derived-Peptide interaction as a novel and potentially early clinical intervention capable of reversing the impact of age-related musculoskeletal diseases. NCAM1-14.3.3.ζδ-derived-Peptide interaction demonstrated efficacy across diverse age-related conditions characterized by vascular and bone loss, including osteoporosis, osteoarthritis and rheumatoid arthritis, underscores its promising potential as a therapeutic target for a spectrum of bone-related disorders.

## Discussion

In this ground-breaking study, employing cell-specific mouse genetics coupled with high-resolution imaging techniques, we not only reveal but also illuminate with unprecedented clarity a hitherto undiscovered and remarkably potent anabolic pathway. The 14.3.3.ζδ-derived-Peptide-NCAM1 interaction emerges as the driving force behind the robust expansion of highly angiogenic Clec14a+ endothelial cells, playing a pivotal role in promoting bone osteogenesis and genesis of new bone.

The robustness of our findings is underpinned by the meticulous application of inducible cell-specific mouse genetics to delineate the role of highly angiogenic Clec14a+ endothelial cells and the NCAM1-14.3.3.ζδ-derived-Peptide interaction in these cells. Furthermore, the use of osteoblast-specific transgenics has allowed us to reveal the role of the NCAM1-14.3.3.ζδ-derived-Peptide interaction, creating a nexus between angiogenesis and osteogenesis.

The direct impact of the 14.3.3.ζδ-derived-Peptide-NCAM1 interaction in osteoblasts is unveiled, elucidating its capacity to drive osteoblast differentiation and, consequently, laying the foundation for the genesis of new bone. Mechanistically, in osteoblasts, the NCAM1-14.3.3.ζδ-derived-Peptide interaction operates through the recognized NCAM1 downstream pathway involving β-catenin signalling (Bonfim et al., 2016; He et al., 2023). Moreover, NCAM1-14.3.3.ζδ-derived-Peptide interaction results in suppression of bone resorption. This nexus between NCAM1-14.3.3.ζδ-derived-Peptide interaction in endothelium proves to be instrumental in reversing bone loss in age-related vascular and bone loss diseases. This positions the 14.3.3.ζδ-derived-Peptide-NCAM1 anabolic interaction as a promising and innovative therapeutic target for managing conditions characterized by vascular loss and, consequently, bone loss.

Taken together, these findings lay the groundwork for a transformative approach in therapeutic interventions. The NCAM1-14.3.3.ζδ-derived-Peptide anabolic interaction, with its remarkable capacity to drive angiogenesis and bone formation, emerges as a powerful tool in our arsenal to combat conditions marked by vascular and bone insufficiencies, offering hope for a paradigm shift in the management of age-associated and bone-related disorders. Anabolic endogenous NCAM1-14.3.3.ζδ-derived-Peptide interaction positions it as a viable candidate for preventing and treating bone loss and fracture. Further studies are urgently required to ascertain and harness the therapeutic potential of NCAM1-14.3.3.ζδ-derived-Peptide pathway in the clinical management of patients with excessive bone loss.

## Methods

### Mice

The study employed C57BL/6 mice from Charles River as wild-type subjects for all analyses, unless otherwise specified. The mice were categorized into three age groups: juvenile (1-4 weeks), adult (12-16 weeks), and aged (>65 weeks). Both male and female mice were included in the study, and details regarding transgenic mouse lines can be found in the resources table. In the drug treatment phase, mice were randomly assigned to receive treatment, while littermates were utilized as sham controls. Mice underwent daily intra-peritoneal (IP) injections over a 10-day period. The injections consisted of either a vehicle control (PBS) or 14.3.3.ζδ-derived-Peptide SVTEQGAELSNEER (400μg). Calcein injections (20mg/kg) were administered on days 8 and 12. The study adhered to the Principles of Laboratory Animal Care outlined by the National Society for Medical Research and the Guide for the Care and Use of Laboratory Animals (National Academies Press, 2011). All Experiments through out the study involving animals were performed according to institutional guidelines and laws, following protocols approved by local animal ethics committees in the UK and in China.

### Tamoxifen treatment for inducible gene deletion

Tamoxifen treatment for inducible gene deletion and genetic lineage tracing was conducted following established protocols (Kusumbe et al., 2014). Tamoxifen (Sigma-Aldrich, T5648) was freshly prepared by initially dissolving it in 100% ethanol and subsequently suspending it in corn oil to achieve a final concentration of 5 mg/ml. For tamoxifen-induced genetic lineage tracing, Cre ERT2 mouse lines, as specified in the figure legends, were employed. To activate Cre recombinase, tamoxifen was administered orally at a dose of 50 mg/kg for three consecutive days. Consistent with prior studies (Kusumbe et al., 2014; Kusumbe et al., 2016), tamoxifen injections were performed in both Cre+ and Cre-mice across all experiments in this study. At the designated time points, mice were euthanized via CO2 asphyxiation, and tissues were collected for subsequent analysis. This methodology adhered to the established procedures outlined in previous publications (Biswas et al., 2023).

### Generation of Clec14a Cre ERT2 mouse line

The mouse Clec14a gene (NCBI Reference Sequence: NM_025809.5) is located on mouse chromosome 12. 1 exon have been identified, with the ATG start codon in exon 1 and TGA stop codon in exon 1. A BAC containing the upstream regulatory sequence of the mouse Clec14a gene was identified. The “CreERT2-polyA” cassette was placed upstream of the ATG start codon in the BAC. The PiggyBac ITRs were inserted into the BAC backbone flanking the genomic insert to facilitate transposition mediated BAC integration. The modified BAC was co-injected with transposase into single-cell stage fertilized eggs from C57BL/6J mice. The pups were genotyped by PCR for the presence of the modified BAC. The positive founder mice were counter screened for transposition. This Clec14a Cre ERT2 mouse line was generated by using the service from Taconic Biosciences.

### Preparation of samples for Immunohistochemistry

To prepare samples for immunohistochemistry, bones were dissected and placed in ice-cold 2% paraformaldehyde (PFA) in PBS, where they were left on ice for 4 hours. Subsequently, the bones underwent processing following established methods (Kusumbe et al., 2015). After rinsing in PBS, bone samples were immersed in 0.5M EDTA (pH 7.4) for a minimum of 36 hours. Following this, they were dehydrated in a solution containing 20% sucrose (Sigma-Aldrich, S9378) and 2% polyvinylpyrrolidone (PVP, Sigma-Aldrich, PVP360) for 48 hours. The prepared bones were then embedded in a mixture of 20% sucrose, 2% polyvinylpyrrolidone, and 8% gelatin (Sigma-Aldrich, G2625). Sectioning of the samples was accomplished at a thickness of 100 µm using a Leica CM3050 cryostat equipped with low-profile blades (Leica, 14035838382). Subsequently, the sections were air-dried before being placed in the freezer for storage. This sample preparation procedure adhered to the methodology previously described previously (Kusumbe et al., 2015).

### Immunostaining of thick bone slices

In the immunostaining process for thick bone slices, meticulous steps were undertaken to ensure precision and reproducibility (Kusumbe et al., 2015). Initially, bone sections underwent a 15-minute air-drying phase followed by a 5-minute rehydration in PBS to optimize the subsequent staining procedure. To enhance permeability, sections were treated with 0.3% Triton X-100 for 10 minutes and subsequently blocked in 5% donkey serum at room temperature (RT). Primary antibodies, critical for target-specific binding, were incubated with the samples after being judiciously diluted in blocking buffer during an overnight incubation at 4°C. The comprehensive list of primary antibodies is thoughtfully documented in the key resources table. To eliminate excess primary antibodies and enhance specificity, samples were subjected to multiple 5-minute washes at RT using PBS. Subsequently, the samples underwent a 2-hour incubation at RT with Alexa fluor-conjugated secondary antibodies, detailed in the key resources table, and the nuclear marker TOPRO-3 or 4’,6-Diamidino-2-Phenylindole (DAPI) at a 1:1000 dilution. Following additional PBS washes, the stained samples were carefully mounted with glass coverslips using Fluoromount-G (Invitrogen, 00-4958-02). To validate specificity, negative controls were implemented, involving staining without primary antibodies and exclusively with secondary antibodies. This detailed protocol ensures clarity and methodological transparency in the immunostaining process, allowing for precise analysis of thick bone slices.

### Microscope set-up and Imaging acquisition

The confocal imaging setup and image acquisition process were conducted with precision and detail using the Zeiss Laser Scanning Confocal Microscope LSM880. Z-stacks of immunostained sections were captured utilizing both the 20X Plan Apo/0.8 dry lens and the 10X Plan Apo 0.45 WD=2.0 M27 dry lens. The imaging configuration featured a Zeiss laser scanning microscope 880 equipped with Axio Examiner, incorporating laser lines at 405, 453, 488, 514, 561, 594, and 633 nm. The system included a Colibri 7 epifluorescence light source with LED illumination, four objectives, a fast-scanning stage with PIEZO XY, a 32-channel gallium arsenide phosphide detector (GaAsP) PMT (photomultiplier tube) in addition to a two-channel standard PMT. The acquisition and analysis software encompassed various functionalities, such as measurement, multichannel, panorama, extended manual focus, image analysis, time-lapse, Z Stack, extended focus, autofocus, and incorporated additional modules, including Experiment Designer and Tiles and Position.

To capture large regions within the thick sections, the tile scan function was employed, and images with a 10% overlap were stitched together using Zen Black (version 3.1, Zeiss) software. To enhance the visualization of organ boundaries, autofluorescence from the 405 channel was converted into greyscale, and the resultant 30% opaque image was manually overlaid with the corresponding TIFF file generated from Imaris. Image generation, analysis, and compilation were carried out using Imaris, Adobe Photoshop, and Adobe Illustrator software, ensuring a comprehensive and sophisticated approach to confocal imaging and subsequent data processing.

For Imaging with the Leica thunder microscope of immunofluorescent-stained bone sections, the Leica DM6 B Thunder Microscope equipped with LAS X software (Version 3.7.4.23463, Leica) was utilized. The microscope configuration included two HC PL FLUOTAR dry objectives (10X and 20X magnification), one HC PL APO 100X oil objective, a Leica K5 camera for fluorescent imaging, Leica CTR6 LED power supply, Leica EL6000 (120 W metal halide) external light source, and a diode-pumped, solid-state laser (355 nm) with 7 cube filters (LMD-BGR, LMD-GFP band pass, LMD-GFP long pass, LMD-Cy3, LMD-DAPI, LMD-Alexa594, LMD-CFP, LMD-GFP/Cy3, LMD-YFP, LMD-Cy5). Z-stacks were acquired using Leica SmartMove Controls for z (focus) movement and x, y (stage) movement for focusing. The LAS X software facilitated image acquisition, offering tools for adjusting exposure times, contrast, and brightness, Live-imaging-mode, a memory function for two z-positions, tile scan, a subsequent merging function, and Thunder Computational Clearing Software.

During imaging, bone sections were inspected in Live mode using appropriate channels selected based on the fluorescence spectrum of the respective secondary antibody to identify a region of interest. Exposure time, brightness, and contrast were adjusted for optimal visualization of the brightest sample in the staining panel. In instances of high background signal, exposure was reduced to mitigate unspecific signals from endogenous background structures. Exposure settings were standardized for subsequent samples within an experiment group to ensure comparability of mean fluorescence between samples. Images were captured using the tile scan function, and the resulting tile scans were merged and processed with Thunder Computational Clearing software to enhance image contrast and reduce blurriness. Thunder-processed images were further analysed using Imaris software (version 9.6.0, Bitplane) for subsequent processing and detailed analysis.

### RNA isolation and quantitative PCR

For the isolation of total RNA from whole bones, the RNeasy Mini Kit (QIAGEN) manufacturer’s protocol was strictly followed. Crushed bones, prepared using a mortar and pestle according to established procedures (Kusumbe et al., 2014), underwent subsequent digestion with a mixture comprising 0.2% collagenase IV, dispase (1.25 U/ml) (Thermo Fisher Scientific, catalog no. 171055-041), and DNase I (7.5 mg/ml) (Sigma-Aldrich, catalog no. D4527-10KU) for 45 minutes at 37°C. This digestion step was crucial for effective RNA isolation.

To generate cDNA, 100 ng of isolated RNA per reaction was utilized with the iScript cDNA Synthesis System (Bio-Rad). The resulting cDNA served as the template for quantitative polymerase chain reaction (qPCR) on the ABI PRISM 7900HT Sequence Detection System, using TaqMan gene expression assays. FAM-conjugated TaqMan probes, along with the TaqMan Gene Expression Master Mix (Applied Biosystems, 4369510), were employed for qPCR. The gene expression assays were normalized using endogenous VIC-conjugated Actb probes as standards. RNA samples were promptly processed for complementary DNA preparation using the SuperScript IV First-Strand Synthesis System (Invitrogen, 18091200).

### Micro-CT analysis and histomorphometry

Tibiae were collected; the attached soft tissue in the bone was removed thoroughly and fixed in 2% paraformaldehyde. The fixed tibiae were analyzed using micro-CT (µCT 100). Following scan settings were used Voxel size 6.0 μm, FOV 10.236 mm, Image matrix 1706 x 1706 x 550, Slices 550, Scanned region 3.3 mm, X-ray voltage 70 kVp, and Intensity 85 μA, 6 W. Calcein double labeling was performed to calculate Bone Formation Rate (BFR) and mineral apposition rate (MAR). Mice were given intraperitoneal injections of 10 mg/kg calcein (Sigma, C0875) dissolved in 2% sodium bicarbonate solution on the 10^th^ day and 3^rd^ day before euthanasia. Bones were fixed in 4% PFA, embedded in 8% gelatin and 2% PVP and cryosectioned. Single plane images were acquired from the sections. Sections were stained with von Kossa method to assess mineralized bone. Representative images show cortical bone (diaphysis about 3 mm proximally from the growth plate). Mineral Apposition Rates were calculated from both cortical and trabecular bones.

its (R&D Systems) according to the manual provided by the manufacturer.

### Bone Metastasis

For experimental metastasis assays, MDA-MB-231 cells or PC3 (1 x 10^5^ cells in 100 µl PBS) were injected into the left cardiac ventricle of NOD-SCID mice with a 26½ gauge needle. MDA-MB-231 cells were always injected into female NOD-SCID mice. Successful injection was characterized by pumping arterial blood into the syringe. Mice were treated after 15 days after intra-cardiac injections with 14.3.3.ζδ-derived-Peptide.

### Osteoporosis, Osteoarthritis and Collagen Induced Arthritis mouse models

Collagen-Induced Arthritis (CIA) was used as the experimental model for rheumatoid arthritis. The mice utilized in the experiments were housed under stringent pathogen-free conditions, ensuring a controlled environment, with ad libitum access to food and water. The induction of arthritis involved the subcutaneous injection of 200 μg of bovine type II collagen in complete Freund’s adjuvant (CFA) at the base of the tail in DBA/1 mice, following established protocols (Choudhary et al., 2018).

Destabilization of the medial meniscus (DMM) model of osteoarthritis was used. For the surgical destabilization of the DMM model male mice underwent the procedure as described previously (Glasson et al., 2007). Treatment with 14.3.3.ζδ-derived-Peptide was initiated 12 weeks post-surgery. Ovariectomy was performed induce osteoporosis. 16-weeks-old female C57Bl/6J underwent ovariectomy as previously described. 2 weeks later, mice were culled to acquire baseline measurements or 14.3.3.ζδ-derived-Peptide treated was initiated.

### Single cell data analysis

The count data used in this study was extracted from the openly accessible dataset GEO GSE122467 (https://www.ncbi.nlm.nih.gov/geo/query/acc.cgi?acc=GSE122467) under the accession numbers (GSM3466897, GSM3466898, GSM3466899, GSM3466900, GSM3466901, GSM3937216, GSM3937217) (Kurtova, Antonina V., et al. 2023). Subsequently, the data underwent consolidation and additional processing using the Seurat CRAN package (version 4.0.1) (https://satijalab.org/seurat/) (Satija et al., 2015). Specifically, cells positive for PECAM1 were selectively filtered for subsequent analysis. Following this filtering step, a total of 763 cells remained for the ensuing analysis. The filtered cell data underwent normalization, scaling, and identification of variable genes using SCTransform (Hafemeister & Satija, 2019). Cell clusters were determined using the FindClusters function of the Seurat package with a resolution of 0.4. The resulting cell clustering was visualized using Uniform Manifold Approximation and Projection for Dimension Reduction (umap). Additional visualizations, including umap gene expression overlays, feature plots, and violin plots illustrating cell type-specific marker genes, were generated using Seurat-specific functions.

Further, we used human bone marrow data with accession number E-MTAB-11560 (Biswas et al., 2023) and explore gene expression of PECAM1, CLEC14A, NCAM1, CDH5, EMCN.

### NCAM-1-14.3.3.ζδ-derived-Peptide Protein complex prediction with AlphaFold-Multimer

Neural cell adhesion molecule 1 (NCAM-1) protein sequence obtained from Uniprot (H7BYX6) sequences were run alongside 14.3.3.ζδ-derived-Peptide (SVTEQGAELSNEER) to predict 14.3.3.ζδ-derived-Peptide-potential binding partner interactions using AlphaFold2_multimer (Guo et al., 2022). This analysis was conducted in Google Colab (Shinkai & Itoga, 2022), resulting in the generation of five distinct models through AlphaFold2_multimer. The selection of the most reliable models was based on the predicted local distance difference test (pLDDT), choosing those with the highest predicted confidence. Subsequently, ChimeraX (Goddard et al., 2018; Schaefer et al., 2021) was employed to construct a 3-D model representing the interaction between NCAM-1 and 14.3.3.ζδ-derived-Peptide.

### Statistical analysis and reproducibility

Statistical analysis and reproducibility were integral aspects of our study, with panels typically representing data from multiple independent experiments conducted on different days and involving distinct sets of mice. All the data presented in this study is derived from minimum of three independent experiments. All the sample numbers show biological replicates. It is important to note that sample sizes were not predetermined based on statistical power calculations. Mice were randomly allocated to experiments, and sample processing occurred in an arbitrary order. While blinding techniques were not employed, no animals were excluded from the analyses. Variability within the data was consistently indicated using standard deviation. The presentation of results involved displaying data as mean ± SEM or SD, as appropriate. To assess the statistical significance of differences between two groups, we primarily utilized two-tailed unpaired Student’s t-tests. For comparisons among more than two groups, one-way ANOVAs with Tukey’s multiple comparisons tests were performed. A significance level of P < 0.05 was considered statistically significant. Notations included “ns” for nonsignificant (P > 0.05), “\**” for P < 0.05, “**” for P < 0.01, “\*\*\**” for P < 0.001, and “****” for P < 0.0001. All statistical analyses were conducted using GraphPad with Prism10 (version 10), following its Statistics Guide. It’s important to note that no statistical analysis was used to determine the sample size in our experiments. This comprehensive approach to statistical analysis and reporting ensures transparency and provides a clear understanding of the rigor employed in the interpretation of the experimental results.

## Author Information

The authors do not declare competing financial interests.

## Acknowledgments

This work is funded by Medical Research Council, Career Development Award (MR/P02209X/1), European Research Council Starting Grant (metaNiche, 805201), Leukemia UK (2017/JGF/001), The Royal Society (RG170326), Kennedy Trust for Rheumatology Research Senior Research Fellowship (KENN 15 16 09) and John Fell Fund OUP Research Fund (161/061) to APK.

## Legends to Figure Supplements

**Figure 1– Figure Supplement 1 - Changes in bones after 14.3.3.ζδ-derived-Peptide treatment**

**(A)** Quantitative analysis of relative bone volume (Rel. BV/TV, left panel) and cortical thickness (Ct. Th, right panel) of tibial bones from young mice treated with altered peptide compared to Vehicle (*n* =7).

**(B)** 3D images of tibial bone from young mice treated with 14.3.3.ζδ-derived-Peptide compared to Vehicle; showing adipocytes, labeled with Perilipin (green), and DAPI (blue).

**Data Information:** Two-tailed unpaired t tests (A). nsp > 0.05, **p < 0.01, ***p < 0.001, and ****p < 0.0001. Scale bar: yellow, 300 μm. n represents biological replicates.

**Figure 2 – Figure Supplement 1 Generation of CLEC14a CreER^T2^ transgenics and genetic depletion of CLEC14a+ endothelial cells**

**(A-B-C-D)** Creation steps of a PiggyBac-on-BAC transgenic line expressing CreERT2 under the control of mouse Clec14a promoter in C57BL/6J mouse.

**(E)** Relative fold mRNA expression of osteogenic genes in Clec14a+ endothelial cells after 14.3.3.ζδ-derived-Peptide treatment (*n* =7).

**(F)** Schematic depicting the experimental design, whereby *Clec14a Cre ER^T2^ X iDTA* mice were not treated or subjected to 14.3.3.ζδ-derived-Peptide treatment prior to bone collection.Quantitative analysis of relative bone volume (Rel. BV/TV, left panel) and cortical thickness (Ct. Th, right panel) of tibial bones from Cre^-^ or Cre^+^, *Clec14a Cre ER^T2^ X iDTA* mice after 14.3.3.ζδ-derived-Peptide treatment (*n* =7).

**Data Information:** Two-tailed unpaired t tests (E and F). nsp > 0.05, **p < 0.01, ***p < 0.001, and ****p < 0.0001. n represents biological replicates.

**Figure 3 – Figure Supplement 1 CDH15 and NCAM1 blocking and analysis of *Ncam1* expression during 14.3.3.ζδ-derived-Peptide treatment**

**(A)** Quantitative analysis of relative bone volume (Rel. BV/TV, left panel) and cortical thickness (Ct. Th, right panel) of tibial bones – vehicle, and 14.3.3.ζδ-derived-Peptide treated or in combination with CDH15 agonistic antibody (n=7).

**(B)** Relative fold mRNA expression of *Ncam1* in osteoblasts and Clec14a+ endothelial cells after 14.3.3.ζδ-derived-Peptide treatment (*n* =7).

**(C)** Quantitative analysis of relative bone volume (Rel. BV, left panel) and cortical thickness (Ct. Th, right panel) of tibial bones – vehicle, and 14.3.3.ζδ-derived-Peptide treated or in combination with NCAM1 antibody (n=7).

**(D)** Quantification of Calcitonin Receptor+ cells and CTX-1 levels from Cre^+^ *Col1a Cre ER^T2^ X Ncam1^fl/fl^* and Cre^+^ *Clec14a Cre ER^T2^ X Ncam1^fl/fl^* mice; 14.3.3.ζδ-derived-Peptide treated mice compared to Vehicle

**Data Information:** Two-tailed unpaired t tests (B and D); One-way ANOVA tests with Tukey’s multiple comparisons tests (A and C). nsp > 0.05, **p < 0.01, ***p < 0.001, and ****p < 0.0001. n represents biological replicates.

**Figure 4 – Figure Supplement 1 Increased angiogenesis markers after 14.3.3.ζδ-derived-Peptide treatment in age related musculoskeletal diseases**

Relative fold mRNA expression of *Ncam1* and *Clec14a+* after 14.3.3.ζδ-derived-Peptide treatment in age related musculoskeletal diseases (*n* =6). Two-tailed unpaired t test, ****p < 0.0001. n represents biological replicates.

## Notes

### Competing Interest Statement

The authors have declared no competing interest.

