## Supplementary figures and images for "Stimulation of NCAM1-14.3.3.ζδ-derived Peptide Interaction Fuels Angiogenesis and Osteogenesis in Ageing"

### Supplementary Figure 1

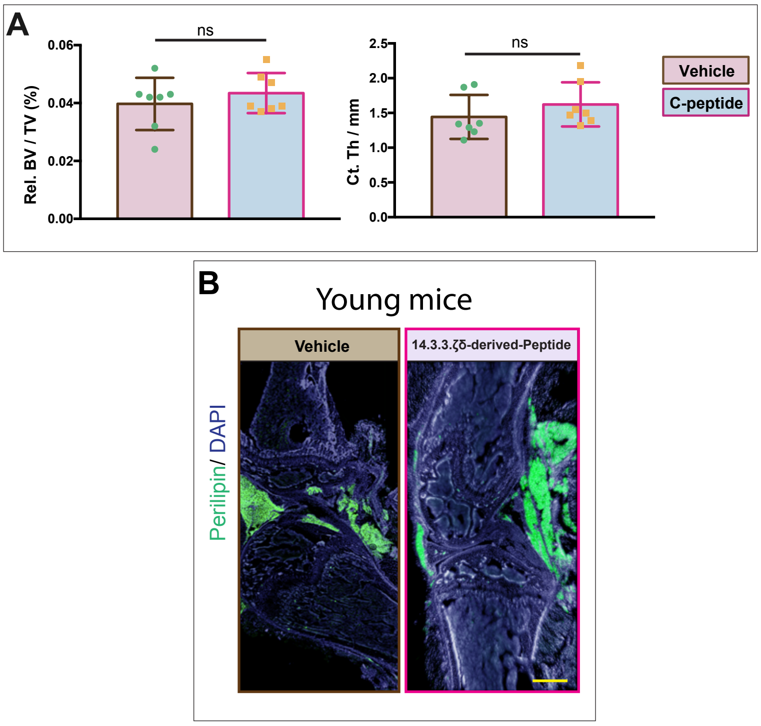


Figure1- figure supplement 1

### Supplementary Figure 2

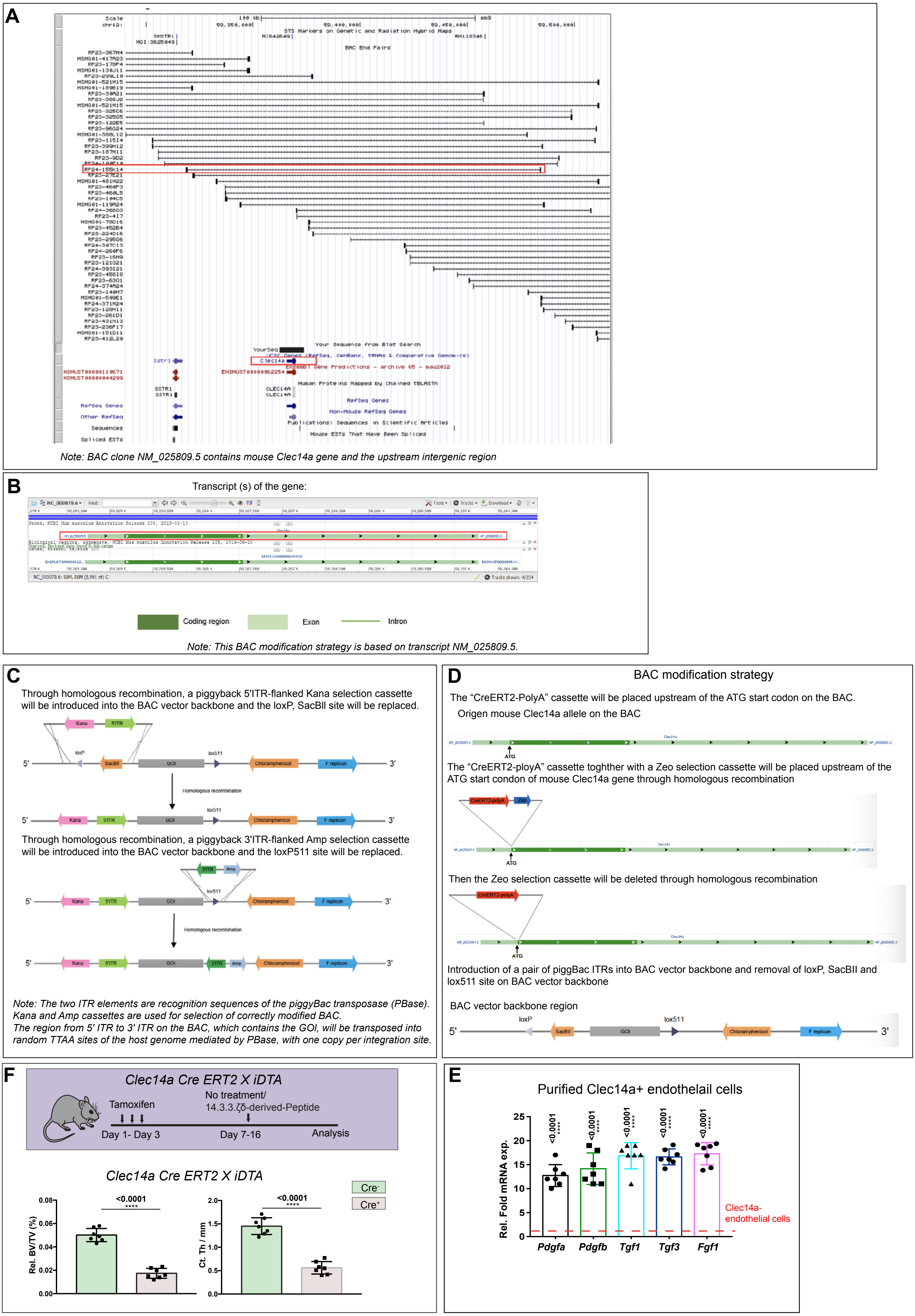


Figure 2 - figure supplement 1

### Supplementary Figure 3

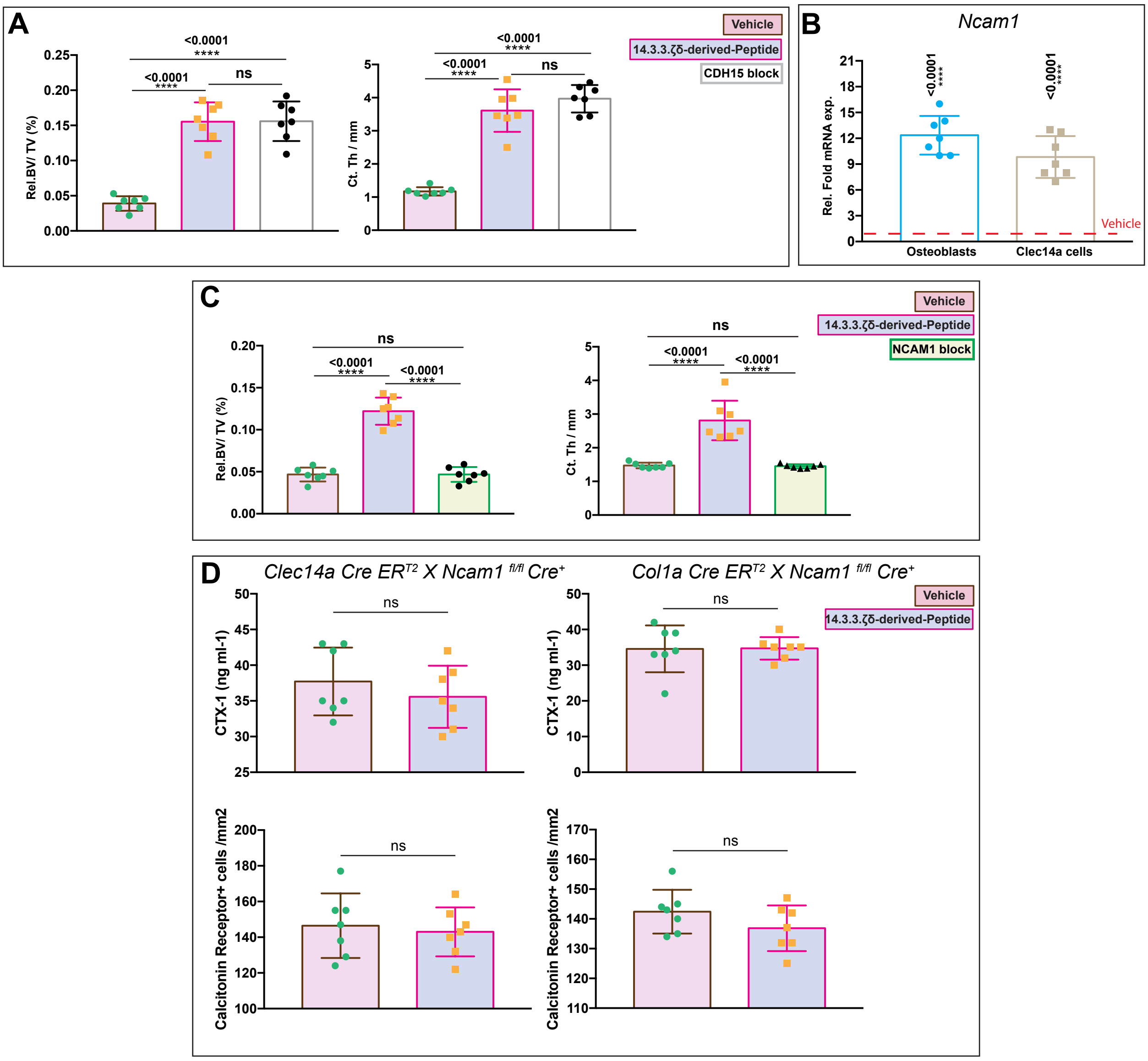


Figure 3—figure supplement 1

### Supplementary Figure 4

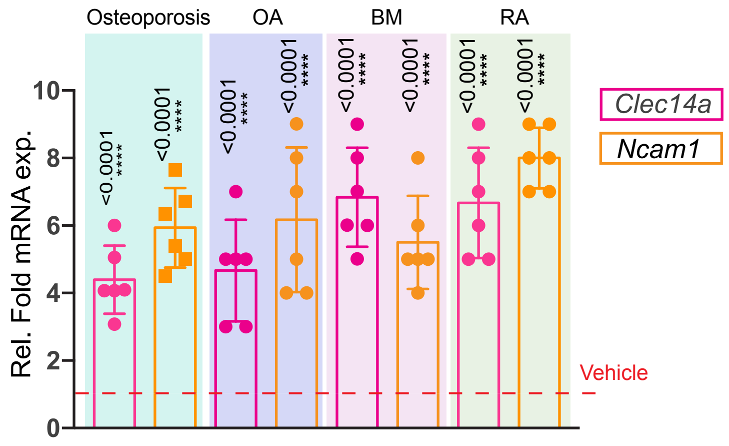


Figure 4—figure supplement 1
